# Reproducible Bioinformatics Project: A community for reproducible bioinformatics analysis pipelines

**DOI:** 10.1101/239947

**Authors:** Neha Kulkarni, Luca Alessandrì, Riccardo Panero, Maddalena Arigoni, Martina Olivero, Francesca Cordero, Marco Beccuti, Raffaele A Calogero

## Abstract

**Background:** Reproducibility of a research is a key element in the modern science and it is mandatory for any industrial application. It represents the ability of replicating an experiment independently by the location and the operator. Therefore, a study can be considered reproducible only if all used data are available and the exploited computational analysis workflow is clearly described. However, today for reproducing a complex bioinformatics analysis, the raw data and a list of tools used in the workflow could be not enough to guarantee the reproducibility of the results obtained. Indeed, different releases of the same tools and/or of the system libraries (exploited by such tools) might lead to sneaky reproducibility issues.

**Results:** To address this challenge, we established the *Reproducible Bioinformatics Project (RBP)*, which is a non-profit and open-source project, whose aim is to provide a schema and an infrastructure, based on docker images and R package, to provide reproducible results in Bioinformatics. One or more Docker images are then defined for a workflow (typically one for each task), while the workflow implementation is handled via R-functions embedded in a package available at github repository. Thus, a bioinformatician participating to the project has firstly to integrate her/his workflow modules into Docker image(s) exploiting an Ubuntu docker image developed ad hoc by RPB to make easier this task. Secondly, the workflow implementation must be realized in R according to an R-skeleton function made available by RPB to guarantee homogeneity and reusability among different RPB functions. Moreover she/he has to provide the R vignette explaining the package functionality together with an example dataset which can be used to improve the user confidence in the workflow utilization.

**Conclusions:** Reproducible Bioinformatics Project provides a general schema and an infrastructure to distribute robust and reproducible workflows. Thus, it guarantees to final users the ability to repeat consistently any analysis independently by the used UNIX-like architecture.

## Background

Recently Baker and Lithgow [1, 2] highlighted the problem of the reproducibility in research. Reproducibility criticality affects to different extent a large portion of the science fields [1]. Since nowadays bioinformatics plays an important role in many biological and medical studies [3], a great effort must be put to make such computational analyses reproducible [4, 5]. Reproducibility issues in bioinformatics might be due to the short half-life of the bioinformatics software, the complexity of the pipelines, the uncontrolled effects induced by changes in the system libraries, the incompleteness or imprecision in workflow description, etc. To deal with reproducibility issues in Bioinformatics Sandve [5] suggested ten good practice rules for the development of a computational workflow (Table 1). A community that fulfill some of the rules suggested by Sandve is Bioconductor [6] project, which provides version control for a large amount of genomics/bioinformatics packages. In this way, old releases of any Bioconductor package are kept available for the users. However, Bioconductor does not cover all the steps of any possible bioinformatics workflow, e.g. in RNAseq wolkflow fastq trimming and alignment steps are generally done using tools not implemented in Bioconductor. BaseSpace [7, 8] and Galaxy [9] represent an example of both commercial and open-source solutions, which partially fulfill Sandve’s roles. Furthermore, the workflows implemented in such environments cannot be heavily customized, e.g. BaseSpace has strict rules for applications submission. Moreover, clouds applications, as BaseSpace, have to cope with legal and ethical issues [10]. On the other hand, Galaxy does not provide standardized metadata to annotate workflows.

**Table 1:** Good practice bioinformatics rules, derived from Sandve et al. [5]

|  |  |
| --- | --- |
| **1** | For Every Result, Keep Track of How It Was Produced |
| **2** | Avoid Manual Data Manipulation Steps |
| **3** | Archive the Exact Versions of All External Programs Used |
| **4** | Version Control All Custom Scripts |
| **5** | Record All Intermediate Results, When Possible in Standardized Formats |
| **6** | For Analyses That Include Randomness, Note Underlying Random Seeds |
| **7** | Always Store Raw Data behind Plots |
| **8** | Generate Hierarchical Analysis Output, Allowing Layers of Increasing Detail to Be Inspected |
| **9** | Connect Textual Statements to Underlying Results |
| **10** | Provide Public Access to Scripts, Runs, and Results |

Recently container technology, a lightweight OS-level virtualization, was explored in the area of Bioinformatics to make easier the distribution, the utilization and the maintenance of bioinformatics software [11–13]. Indeed, since applications and their dependencies are packaged together in the container image, the users have not to download and install all the dependencies required by an application, thus avoiding all the cases where the dependencies are not well documented or not available at all. Moreover, problems related to versions conflicts or updates of the system libraries do not occur, because the containers are isolated from the rest of the operating system.

Among the available container platforms, Docker (http://www.docker.com) is becoming *de facto* the standard environment to quickly compose, create, deploy, scale and oversee containerized applications under Linux. Its strengths are the high degree of portability, which allows users to register and share containers over various hosts in private and public repositories; a more effective resource use and a faster deployment compared with other software.

Although, Menegidio [13], da Veiga [11] and Kim [12] provided a large collection of bioinformatics instruments based on Docker technology, today we are missing a community delivering to bioinformaticians a controlled, but flexible framework to distribute Docker based workflows under the umbrella of a reproducibility framework. Here, we describe the implementation of the Reproducible Bioinformatics Project (RBP, http://reproducible-bioinformatics.org/), aiming to distribute to the bioinformatics community docker-based applications under the reproducibility framework proposed by Sandve [5]. RBP accepts simple docker implementations of *bioinformatics software* (e.g. a docker embedding bwa aligner tool), implementation of *complex pipelines* involving the use of multiple dockers images (e.g. a RNAseq workflow providing all the steps for an analysis starting from the quality control of the fastq to differential expression), as well as *demonstrative workflows* (i.e. docker images embedding the full bioinformatics workflow used in a publication) intended to provide the ability to reproduce published data.

## Implementation

The Reproducible Bioinformatics Project (RBP) reference web page is reproducible-bioinformatics.org. The project is based on three modules (Figure 1): (i) *docker4seq R package* (https://github.com/kendomaniac/docker4seq), (ii) *dockers images* (https://hub.docker.com/u/repbioinfo/), and (iii) *4SeqGUI* (https://github.com/mbeccuti/4SeqGUI).

**Figure 1:**
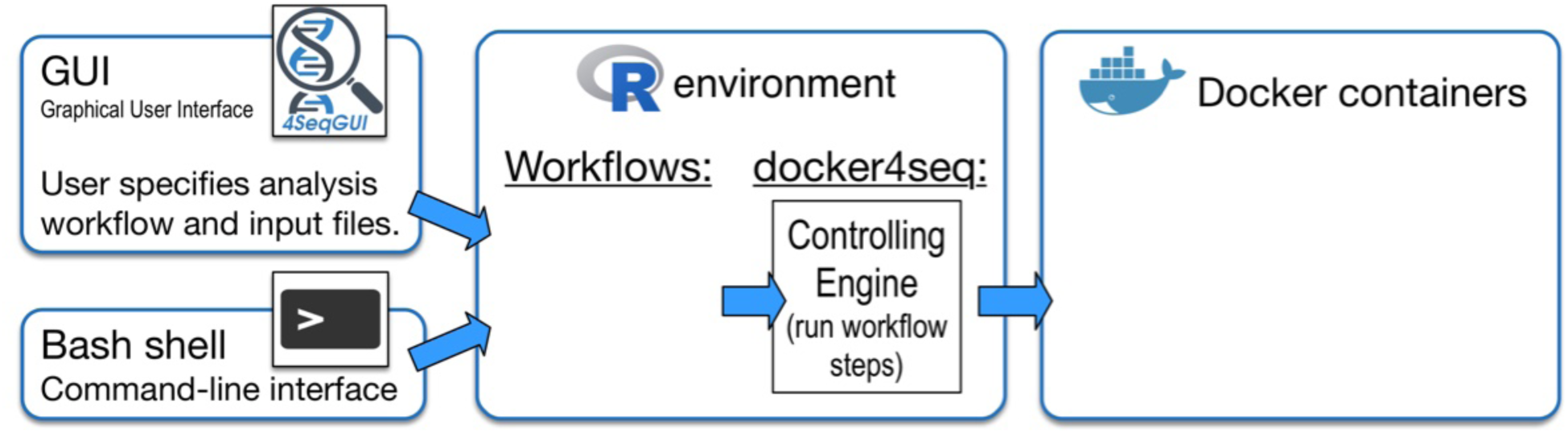
Reproducible Bioinformatics Project structure.

*Docker4seq* package provides the connection between users and docker containers. *Docker4seq* is organized in two branches: stable and development. The transition between development and stable branch is done when a module (R function(s)/docker container(s)) fulfills the 10 rules suggested by Sandve [5] for good bioinformatics practice (Table 1):

The function *skeleton.R* in docker4seq provides a prototype to build a docker controlling function. Acknowledgments of the developer work is provided within the structure of the *skeleton.R*. In *skeleton.R* there is a field indicating developer affiliation and email for contacts. In docker images repository *docker.io/repbioinfo* is available an Ubuntu image, as prototype for the creation of a docker image compliant with the RBP specifications. Developer is free to decide to use this prototype or to adapt a different Linux docker distribution for his/her application. Docker images designed by the core developers of RBP are located in *docker.io/repbioinfo* (docker.com), the images developed by third parties can be instead placed in any public-access docker repository. RBP requires that any operation, implying the use of any R/Bioconductor packages or the use of an external software, has to be implemented in a docker container. Only reformatting actions, e.g. table assembly, data reordering, etc., can be handled outside a docker image.

Any new RBP module (R function(s)/docker image(s)) must be associated with an explanatory vignette, accessible online as html document, and to a set of test data, also accessible online. Thus, all instruments needed to acquire confidence on module functionalities are provided to the final user.

Docker images are labelled with the extension YYYY.NN, where YYYY is the year of insertion in the stable version and NN a progressive number. YYYY changes only if any update on the program(s), implemented in the docker image, is done. This because any of such updates will affect the reproducibility of the workflow. Previous version(s) will be also available in the repository. NN refers to changes in the docker image, which do not affect the reproducibility of the workflow. A new module can be submitted to the and RBP core team will verify the compliance with Sandve [5] rules. Ones validated, the R functions controlling the new module are inserted in *docker4seq* stable release. Partially validated modules will be placed in development branch and moved to stable one when compliance with Sandve’s rules is fulfilled. 4SeqGUI is a Java based graphical interface to docker4seq functions. It is designed to provide a GUI to users having limited knowledge of R scripting. Currently the GUI embeds only general-purpose workflows, such as RNAseq, miRNAseq and Chip-seq workflow.

## Results

The stable branch of *docker4seq R package* contains all the R functions required to handle all the steps of RNAseq workflow (Fig. 2A), ChIPseq workflow (Fig. 2B), and miRNAseq workflow (Fig. 2C). *Docker4seq* also provides a wrapper function for the *bcl2fastq* Illumina tool to convert the Illumina sequencer output in demultiplexed fastq files (Fig. 2). Then, the fastq files can be handled with any of the three different workflows. The counts table produced by RNAseq or miRNAseq workflows can be used for data visualization *(pca*, principal component analysis function), to evaluate the statistical power of the experiment *(experimentPower* function), to define the optimal sample size of the experiment for the detection of differentially expressed genes *(sampleSize* function) and to detect differentially expressed genes/transcripts (*wrapperDeseq2* function). Sample size/statistical power estimation of the experiment and differential expression are calculated respectively via RnaSeqSampleSize [14] and DESeq2 Bioconductor packages [15].

**Figure 2:**
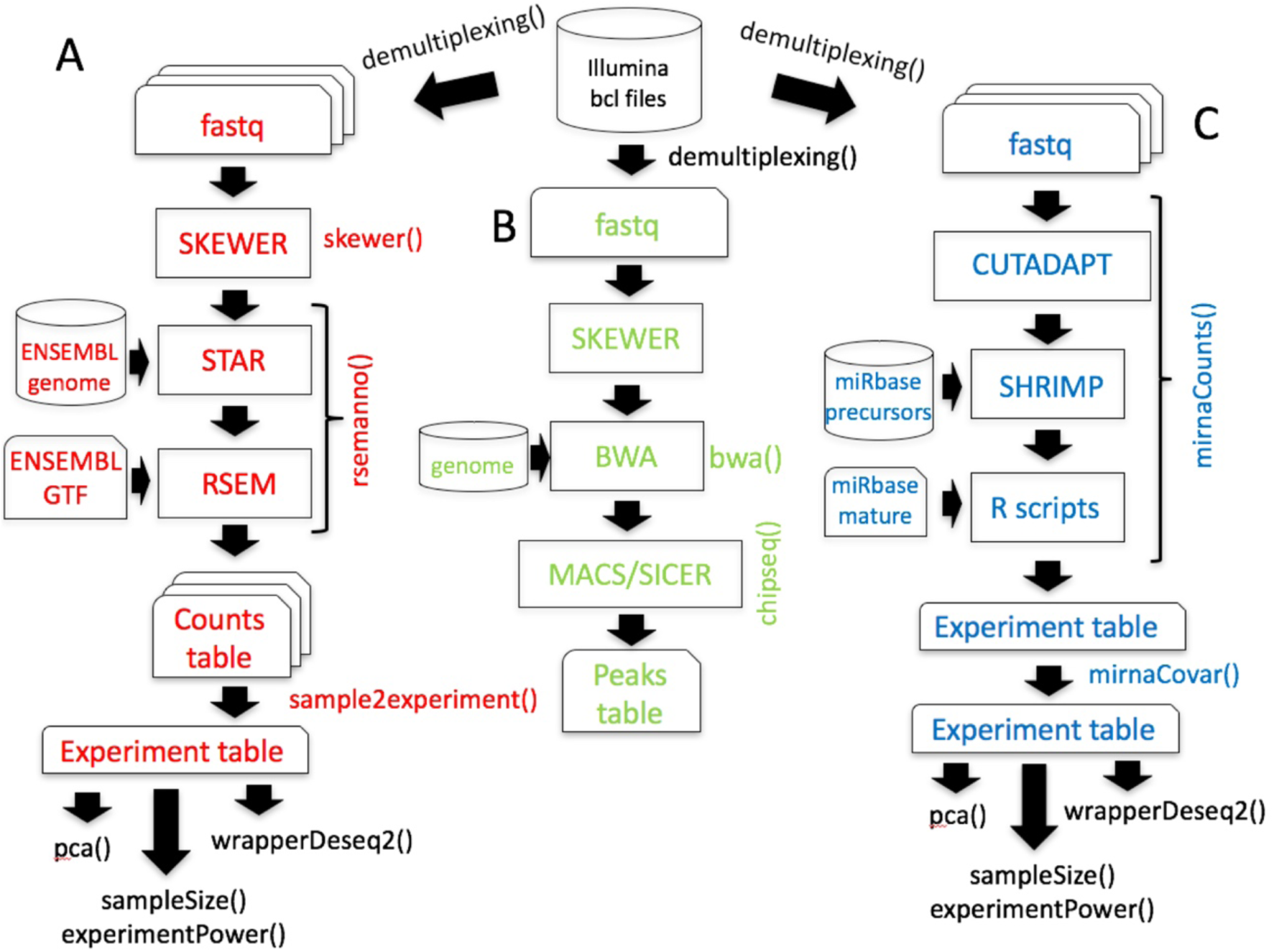
Workflows available in the stable branch of docker4seq. A) Whole transcriptome sequencing workflow, B) ChIP sequencing workflow, and C) miRNA sequencing workflow. The names followed by parenthesis are the docker4seq functions used to execute the analysis steps. Black indicate elements in common among more than one workflow.

In the development branch, the main effort of the core developers is focused in providing workflows for DNA and RNA somatic variant calling. The DNA variant calling workflow embeds the pre-processing procedure suggested by the GATK best practice (Fig. 3A). RNAseq data preparation for variant calling (Fig. 3C) requires the use of STAR 2 step procedure [16], which provides significantly increased sensitivity to novel splice junctions. Then, after sorting and duplicates marking, OPOSSUM [17] is used to remove intronic regions and to merge overlapping reads. We have also implemented a specific procedure (Fig. 3B), based on xenome software [18], to discriminate between human reads and mouse host reads in the sequences produced by the analysis of patients derived xenografts (PDX, [19]). As part of the somatic variant calling workflow we are implementing MUTECT 1 and 2 [20] (Fig. 4A) to call somatic variants as well as PLATYPUS [21] for extracting information of joined-samples SNVs (Fig. 4B).

**Figure 3:**
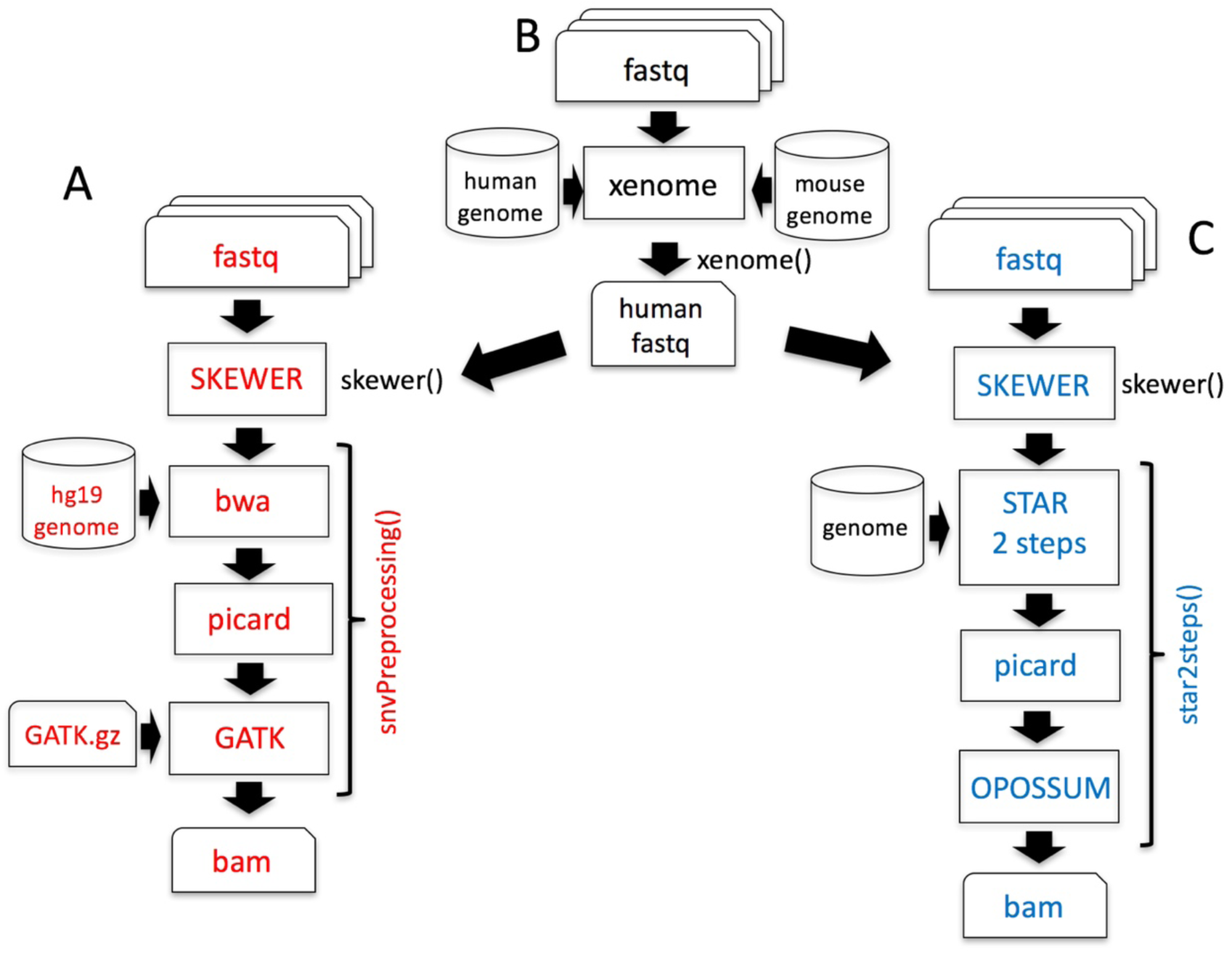
Variant calling workflows under refinement in the development branch of docker4seq. A) SNVs calling in DNA workflow. The function *snvPreprocessing* requires that users provides its own copy of the GATK software, because of Broad Institute license restrictions. This function returns a bam file sorted, with duplicates marked after GATK indel realignment and quality recalibration. B) Data preprocessing for samples derived by Patient Derived Xenografths (PDX). The *xenome* function discriminates between the mouse host reads and the human tumor reads, then DNA or RNA SNV calling workflows can be applied. C) SNVs calling in RNA workflow. The function *star2steps* generates a sorted bam, where duplicates are marked and processed by opossum for removal of intronic regions and merging of overlapping reads. The names followed by parenthesis are the docker4seq functions used to execute the analysis steps. Black indicate elements in common between more than one workflow.

**Figure 4:**
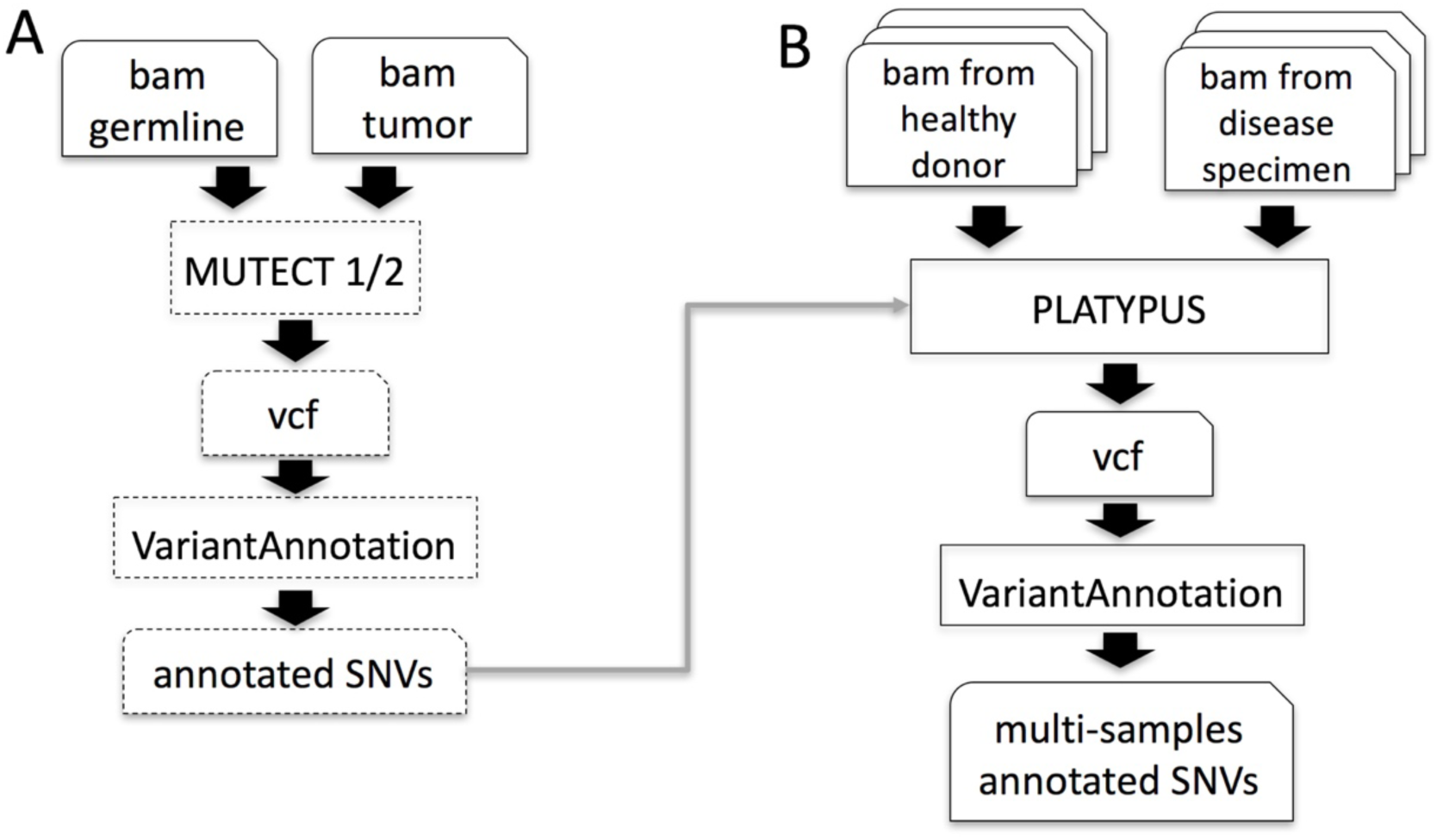
Variant calling workflows under development in the development branch of docker4seq. A) Somatic SNVs detection using GATK MUTECT 1 or 2. B) Platypus based join mutations caller. Dashed blocks are not implemented, yet.

We are also expanding the RNAseq module adding the reference-free Salmon aligner [22], which employs less memory for the alignment task than STAR, but providing similar results [23]. Finally, HashClone framework (Accepted for publication in BMC Bioinformatics), a new suite of bioinformatics tools providing B-cells clonality assessment and minimal residual disease (MRD) monitoring over time from deep sequencing data, was integrated in the *Docker4seq* package. In particular, a parallel version of the standard HashClone workflow (Fig. 5) was developed exploiting the docker architecture.

**Figure 5:**
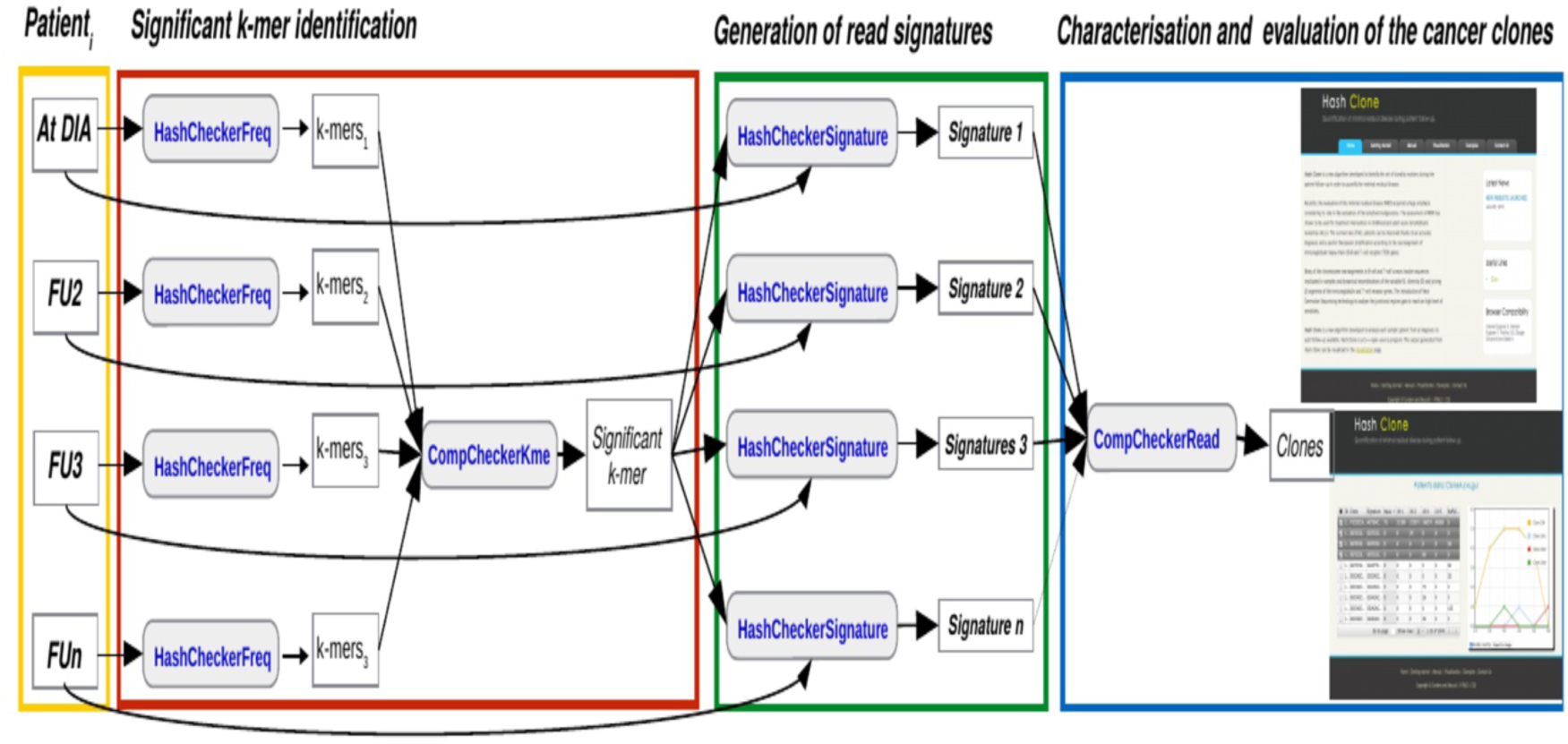
HashClone pipeline. The HashClone strategy is organized in three steps: The first step (red box) is used to detect k-mer in all patients’ samples. The second step (green box) focus on the generation of sequence signatures leading to the identification of the set of putative clones present in each of the patients’ sample; the third step (blue box) is used to the characterization and evaluation of the cancer clones.

All the modules described above are implemented in 18 docker images deposited in the docker hub (https://hub.docker.com/u/repbioinfo/).

As part of the RBP we have also developed a GUI, 4SeqGUI (https://github.com/mbeccuti/4SeqGUI). The GUI is implemented in JAVA and can be exploited to perform whole transcriptome sequencing workflow (Fig. 2A), ChIP sequencing workflow (Fig. 2B), and miRNA sequencing workflow (Fig. 2C).

## Discussion

Bioinformatics workflows are becoming an essential part of many research papers. However, absence of clear and well-defined rules on the code distribution make the results of most published researches unreproducible [24]. Recently, Almugbel and coworkers [25] described an interesting infrastructure to embed Bioconductor based packages. However, Bioconductor does not cover all steps of any possible bioinformatics workflow, thus providing a limited framework for developing complex pipelines. Differently, RBP represents a new instrument, which expands the idea of Almugbel [25], providing a more flexible infrastructure allowing the bioinformatics community to spread their work under the guidance of rules, which guarantee inter-laboratory reproducibility and do not limit docker implementations to Bioconductor packages. RBP core developers created frameworks for RNA/miRNA quantification and analysis. ChIPseq workflow was also developed and variant calling workflows for DNA and RNA are under active development. A peculiar feature of RBP is the acceptance of *demonstrative workflows*, i.e. bioinformatics procedures described in a biological/medical paper. A demonstrative workflow is wrapped in a docker image and it is supported by a tutorial, which describes step by step how the analysis is done to guarantee the reproducibility of published data.

## Availability and requirements

**Project name:** Reproducible Bioinformatics Project

**Project home page:** http://reproducible-bioinformatics.org

**Operating system:** UNIX-like

**Programming language:** R

**Other requirements:** docker version 17.05.0-ce or higher

**License:** GPL.

## Declarations

### Competing interests

None

### Funding

This work has been supported by the EPIGEN FLAG PROJECT

### Authors’ contributions

NK and LA equally contributed to the development of miRNA workflow and all the other tools. RP and FC developed the RNAseq workflow and refined the ChIPseq workflow. MA and MO performed applications testing. MB and RAC developed the rules to submit tools and workflows to the Reproducible Bioinformatics community. RAC and MB equally supervised the overall work.

